# Extracellular Vesicles from Multiple Sclerosis White Matter Exhibit Synaptic, Mitochondrial, Complement, and Aging-related Pathway Dysregulation

**DOI:** 10.1101/2025.06.02.657509

**Authors:** Larissa Jank, Madathiparambil Kumaran Satheesh Kumar, Taekyung Ryu, Rohit Thapa, Olesia Gololobova, Timothy Niepokny, Peter A. Calabresi, Kenneth Witwer, Ranjan Dutta, Chan Hyun Na, Pavan Bhargava

## Abstract

Extracellular vesicles (EVs) are increasingly recognized as mediators of central nervous system (CNS) function and pathologies, including multiple sclerosis (MS). While plasma-derived EVs have been explored as biomarkers in MS, little is known about EVs in CNS tissue. Here, we characterize EVs from postmortem white matter (WM) of MS and control brains. EVs were separated by differential centrifugation followed by size exclusion chromatography and characterized using nanoflow cytometry, single-particle reflectance imaging sensing (SP-IRIS), and transmission electron microscopy. EV size, yield, and morphology did not differ significantly between MS and control samples. Proteomic analyses revealed downregulation of synaptic and mitochondrial proteins and upregulation of complement and inflammatory proteins and pathways in MS WM EVs. This suggests that EVs reflect ongoing synaptic pathology, metabolic dysfunction, and CNS-compartmentalized inflammation and that they may actively contribute to these pathological processes. Deconvolution analyses suggests a shift in EV cellular origin, with an increased astrocytic and decreased neuronal EV contributions in MS. Several proteomic changes we observed in CNS-derived EVs have also been reported in circulating EVs of people with MS, establishing this CNS tissue EV study as a valuable resource for identifying biomarker candidates for brain-derived plasma EV studies.

**GRAPHICAL ABSTRACT:** 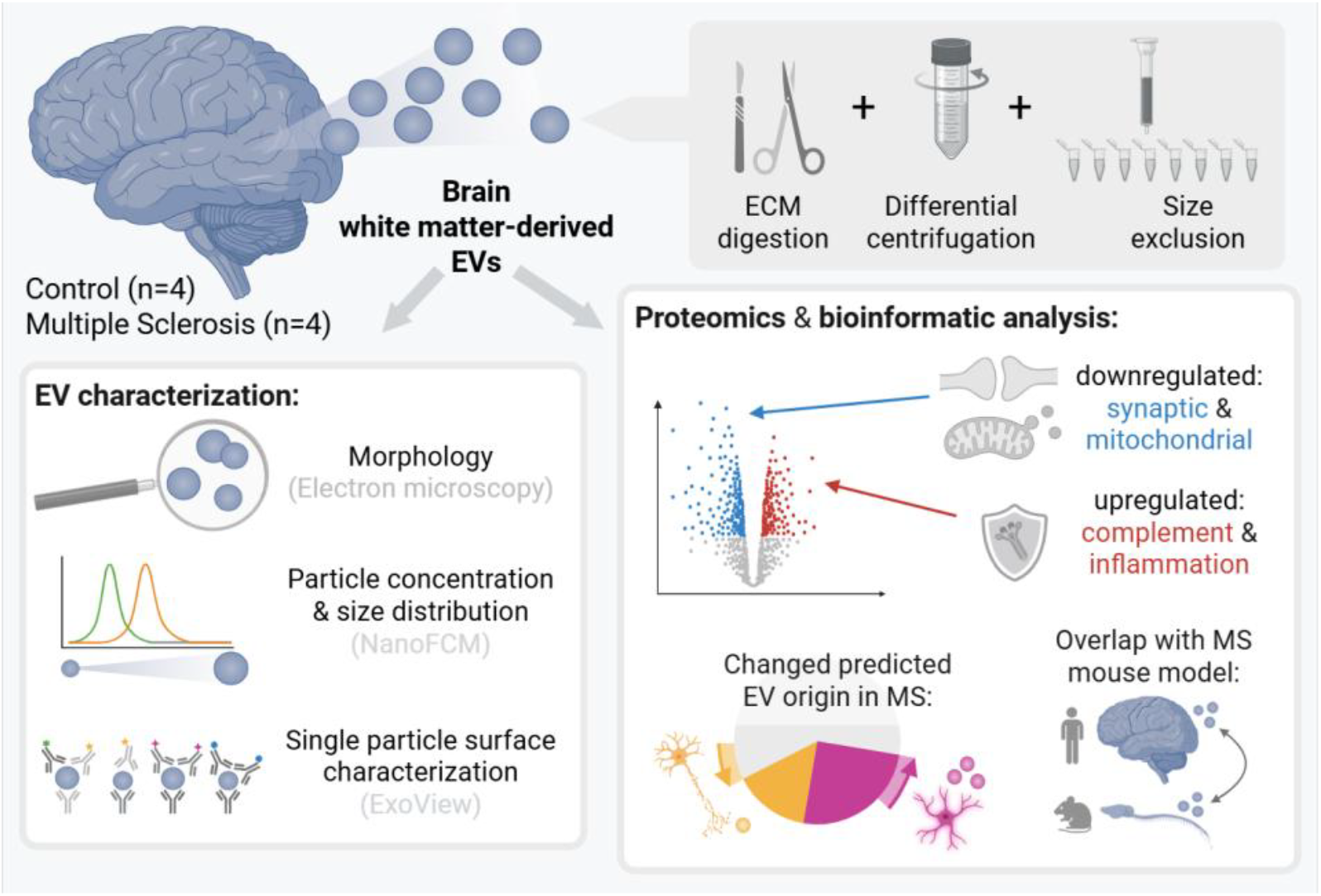

## INTRODUCTION

Extracellular vesicles (EVs) are membrane-delimited particles released by all cells, including those in the central nervous system (CNS). They have a multitude of functions, maintaining homeostasis and promoting healthy CNS function, but can also mediate pathological processes such as neuroinflammation and neurodegeneration *(2)*.

In the field of neuroimmunology – and particularly in multiple sclerosis (MS) research – EVs have emerged as both candidate biomarkers and potential effectors of disease. Recent studies have increasingly focused on using brain-derived EVs from blood plasma to non-invasively monitor CNS-intrinsic neurological disease processes. However, despite the expanding literature on brain-derived EVs in MS, the proteomic landscape of EVs directly isolated from CNS tissue remains poorly characterized. This limits our understanding of how well circulating brain-derived EVs reflect the molecular changes occurring within the brain.

To address this research gap, we separated and profiled EVs from postmortem white matter tissue of people with MS and matched controls. Our findings present a comprehensive bioinformatic analysis of pathways dysregulated in CNS-derived EVs in MS and thereby provide novel insights into the potential role of EVs as mediators of MS pathological processes, including compartmentalized inflammation, synaptic pathology, and mitochondrial dysfunction. Further, this study is a valuable resource to identify new candidates for EV biomarker studies and to align changes observed in brain-derived plasma EV biomarker studies with changes in the CNS.

## METHODS

For a more detailed description of the methods, please refer to the supplementary methods file.

### Tissue Collection and Processing

Postmortem brain tissues from 4 MS patients and 4 matched controls were obtained from the Cleveland Clinic MS brain donation program *(1)* (Patient demographics in Table 1, with further details in Supplementary Table 3). MS patients were diagnosed according to the revised McDonald criteria *(3)*. Cryostat sections (14 μm) were cut from fresh frozen tissue blocks containing subcortical white matter and stained with proteolipid protein (PLP) to determine the presence of demyelinating lesions. Based on the staining results, normal-appearing white matter (myelinated) regions were identified and scored on the frozen block with a scalpel and then collected for EV separation.

**Table 1.** Details of tissues examined in this study.

|  | **Ctrl** | **MS** | *p-value* |
| --- | --- | --- | --- |
| **Age** | 66 ± 9.35 | 54.5 ± 5.5 | 0.12 |
| **Sex** | 2 Female 2 Male | 4 Female |  |
| **Race** | 2 African American,<br>2 Caucasian | 1 African American,<br>3 Caucasian |  |
| **Post-Mortem**<br>**Interval (hrs)** | 15.5 ± 6.10 | 8.44 ± 1.60 | 0.10 |
| **MS subtype** | NA | 1RRMS, 2SPMS,<br>1PPMS |  |
| **Disease Duration**<br>**(yrs)** | NA | 25.25 ± 8.17 |  |
| **Final EDSS** | NA | 8.375 ± 0.65 |  |

### Extracellular Vesicle Separation

EVs were separated using a previously published protocol with slight modifications *(4)*. Briefly, frozen tissue samples were diced on dry ice. A portion was preserved for brain tissue proteomics. The remaining tissue (∼400 mg) was digested in Hibernate-E medium containing 75 U/mL collagenase III for 20 min at 37°C, then immediately chilled and supplemented with protease/phosphatase inhibitors. Samples were then subjected to differential centrifugation (300×g for 10 min, 2,000×g for 15 min, and 10,000×g for 30 min), filtered (0.22 µm), and concentrated to 700 µL. Samples were loaded onto qEV 70 nm columns (IZON), and EV-rich fractions (F7-10) were collected, pelleted at 110,000×g for 90 min, and gently resuspended in 70 µL Dulbecco’s phosphate-buffered saline (DPBS). Aliquots (50 µL for proteomics, 20 µL for other characterization) were stored at -80°C.

### Transmission Electron Microscopy (TEM)

EVs were adsorbed onto glow-discharged carbon-coated copper grids (EMS CF400-CU-UL), washed with TBS and distilled water, and negatively stained with 1% uranyl acetate. Grids were dried and imaged using a Hitachi 7600 TEM. A pooled EV sample (consisting of all four samples) was imaged for each group across eight fields with additional overview images (See Supplementary Figure S1).

### Nanoflow Cytometry

EV size distribution and concentration were measured using a Flow NanoAnalyzer (NanoFCM) following the manufacturer’s protocol. Calibration was performed using 250 nm fluorescent silica particles and a 68–155 nm size standard cocktail. EVs were diluted 1:1000 in DPBS and acquired for 1 min at 1 kPa (1,500–10,000 events/min). Data was processed with the NanoFCM Professional Suite V2.0 software, correcting for dilution and subtracting the background of a DPBS-only control.

### Single-Particle Interferometric Reflectance Imaging Sensor (SP-IRIS)

Tetraspanin profiling was carried out using the ExoView R100 platform with EV-TETRA-C ExoView kits, as previously described *(5)*. EVs (10 µL) were diluted with 35 µL incubation buffer and loaded onto chips for 16h. After thorough washing, chips were stained with fluorescent antibodies targeting CD9, CD63, and Syntenin-1. Chips were imaged using SP-IRIS and analyzed with NanoViewer 2.8.10.

### Proteomic Sample Preparation and LC-MS/MS

Proteomics was performed as previously described *(6)* with minor changes. EVs were lysed in 8 M urea and 50 mM TEAB, reduced with 10 mM TCEP, and alkylated with 40 mM CAA for 1 h. Proteins were digested with Lys-C (1:100, 3 h, 37°C), followed by trypsin digestion (1:50, overnight, 37°C). Peptides were desalted using C18 StageTips, labeled with 18-plex TMTpro reagents, and fractionated into 24 concatenated fractions via high-pH reverse-phase chromatography. Samples were vacuum-dried and reconstituted in 0.5% formic acid. LC-MS/MS was conducted using an Orbitrap Fusion Lumos coupled with an Ultimate 3000 RSLCnano system. Peptides were separated over a 120 min gradient, with MS1/MS2 resolution at 120,000/50,000. HCD was applied at 35% NCE. AGC targets were set to 1 million (MS1) and 50,000 (MS2), with injection times of 50/100 ms. The isolation window was 1.6 Da, and lock mass calibration (m/z 445.12002) was used.

### LC-MS/MS Data analysis

MS/MS data were searched against the UniProt human database using MSFragger via Proteome Discoverer *(7)*. Top 10 peaks per 100 m/z were selected. Parameters included: trypsin digestion with up to 2 missed cleavages, fixed carbamidomethylation on cysteine and TMTpro tags on peptide N-termini and lysine residues, variable methionine oxidation, and MS1/MS2 tolerances of 10/20 ppm. FDR was set at 1% using Percolator. Reporter ion quantification used centroid mode, S/N >10, and both unique/razor peptides. Protein grouping applied strict parsimony, excluding non-unique groups.

### Bioinformatic and Statistical Analysis

UniProt IDs were mapped to Entrez IDs and gene symbols using AnnotationDbi and org.Hs.eg.db *(8, 9)*. PCA was performed using the stats package and visualized with ggplot2 *(10)*. Data were log2-transformed and median-centered. Limma *(11)* was used for differential abundance analysis. Pathway analysis included Gene Set Enrichment Analysis (GSEA) and Over-Representation Analysis (ORA) using ClusterProfiler *(12)* with GO.db *(13)*, KEGG *(14)*, and Reactome *(15)* databases. We used CompareCluster from ClusterProfiler to assess enriched cellular components in EVs compared to tissue, using the top 10% most abundant proteins from tissue and EV samples as the input for this analysis.

Shared/distinct pathways changed in tissue and EVs were identified by ORA of the top up/downregulated (with MS) proteins in each sample type using CompareCluster.

Deconvolution analysis was performed using CIBERSORTx *(16)* with human glial deconvolution matrix *(17, 18)*, and validated with BRETIGEA *(19)*. Limma was used to compare the estimated cell type of origin proportions. Synaptic subsets were explored using GSEA of GO terms related to presynaptic/postsynaptic and excitatory/inhibitory proteins. SynGO *(20)* was used for pathway analysis and sunburst visualization. To assess aging signatures in MS EVs, we performed GSEA with the SenMayo dataset *(21)* and a CSF proteomics aging signature *(22)*, and used a proteomic clock tool based on the CSF dataset. Finally, MS EV profiles were compared to chronic experimental autoimmune encephalomyelitis (EAE) EV proteomic data *(6)*. Protein-wise fold changes were correlated, and shared/distinct pathways were identified by ORA of the top 150 up/downregulated proteins using CompareCluster. All figures were generated in R and finalized in Adobe Illustrator.

## RESULTS

### There is no significant difference in median size, yield, or morphology of NAWM-derived EVs from people with MS compared to controls

EVs were separated from the white matter of four controls (no known neurological disease) and four postmortem human MS brains using differential centrifugation followed by size exclusion chromatography and ultracentrifugation based on a previously published protocol *(4)* (Fig. 1A; sample details in Table 1 with additional information in supplementary table 3). Control and MS EVs were characterized using nanoflow cytometry, ExoView single-particle interferometric reflectance imaging sensor, and transmission electron microscopy (TEM) according to the MISEV criteria *(23)*. There was no significant difference in EV particle yields per tissue mass from MS or control brains. While there was also no significant difference in median particle size, MS brain-derived EVs showed a different distribution pattern in the size histogram and a trend (*p=0*.*08*) towards a higher percentage of particles >80nm compared to control brain-derived EVs (Fig. 1B, supplementary Fig. S1A). Exoview analysis revealed that CD9+ EVs were the most abundant in both MS and Ctrl EVs (supplementary Fig. S1B-C). Overall, the distribution of EV subpopulations was similar in both groups, with the only significant difference being fewer CD63+ CD9+ double-positive particles in MS compared to control EV samples (Fig. 1C). In TEM, EVs from both groups had a typical “cup-shaped” morphology (Fig. 1D, further images in supplementary Fig. S1D). The purity of EVs was further confirmed in cellular component enrichment analysis using the top 10% most abundant proteins in EVs compared to tissue samples. EV samples were enriched for proteins of exocytotic vesicle origin and depleted of proteins of cytoskeletal and mitochondrial origin compared to tissue (Fig. 1E). 88% of the Vesiclepedia *(24)* and 91% of the Exocarta *(25)* top 100 EV proteins were detected in EVs (Fig. 1F).

**Fig. 1.**
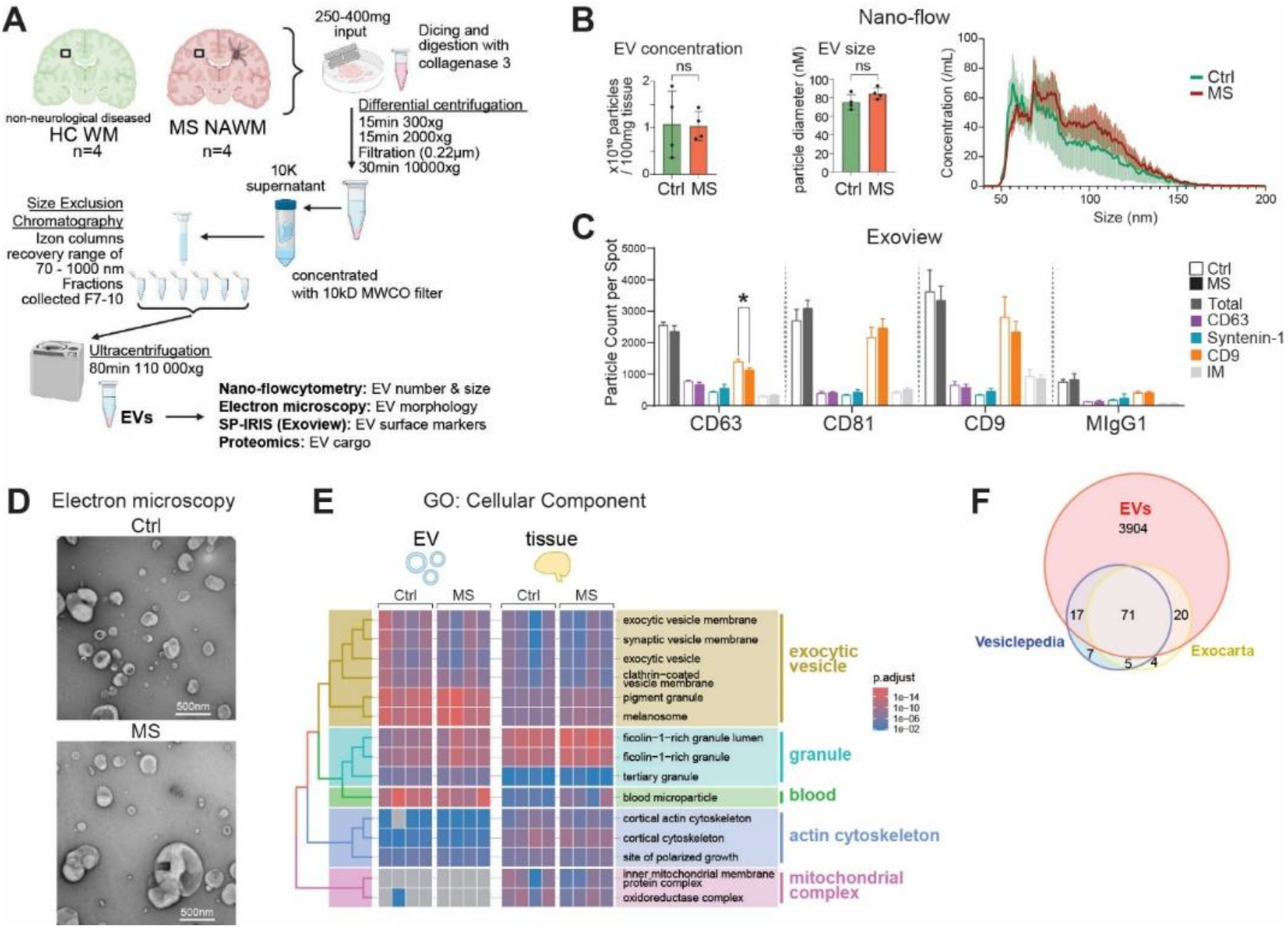
EV separation and characterization. **(A)** Workflow schematic of EV separation using differential centrifugation, size exclusion chromatography, and ultracentrifugation (created using BioRender.com). **(B)** Bar graphs showing the mean particle concentration and median particle size ± SD and a histogram of the full EV size distribution measured using nanoflow cytometry (n = 4 samples for MS NAWM and control [Ctrl] WM EV samples each). Statistical significance was determined using a Student’s t-test (ns = not significant). **(C)** Bar graph (mean particle count ± SD) showing colocalization of tetraspanins on the EV surfaces using ExoView. Statistical significance was determined using a two-way ANOVA with multiple comparisons (* = p<0.05). **(D)** Representative electron microscopy images of negatively stained EVs from control WM and MS NAWM. **(E)** Treeplot and heatmap showing a comparison of the overrepresented GO cellular compartment terms in the EVs and tissue of the MS NAWM and control WM samples. **(F)** Number of the top 100 Vesiclepedia and Exocarta proteins present in the EV proteome.

### Proteomic analysis of WM-EVs reveals changes in inflammatory, synaptic, and mitochondrial proteins, as well as differences in the contribution of astrocytes and neurons to the EV pool in MS EVs compared to controls

Proteomic analysis of the MS and control white matter EVs detected 4029 proteins, of which 291 were differentially abundant (in this exploratory study, defined as *p<0*.*05*). Proteomics of the corresponding tissue detected 7663 proteins, of which 267 were differentially abundant (supplementary tables 1-2). Principal component analysis distinguished EV samples into disease and control groups, despite one control sample (Ctrl4) differing from the remaining controls (Fig. 2A). The top upregulated proteins in MS EVs compared to control EVs included proteins involved in lipid metabolism (ADIRF, FABP5), reactive astrocytosis (CD44, VIM), inflammatory activation (S100A1, A6, A13, CP, FSTL4), and negative regulation of inflammation/cell activation (ANXA1, SZT2). Meanwhile, mitochondrial (LRPPRC, ATP6V0C) and synaptic (SLC6A12, SYT12) proteins were among the top downregulated proteins in MS EVs (Fig. 2B).

**Fig. 2.**
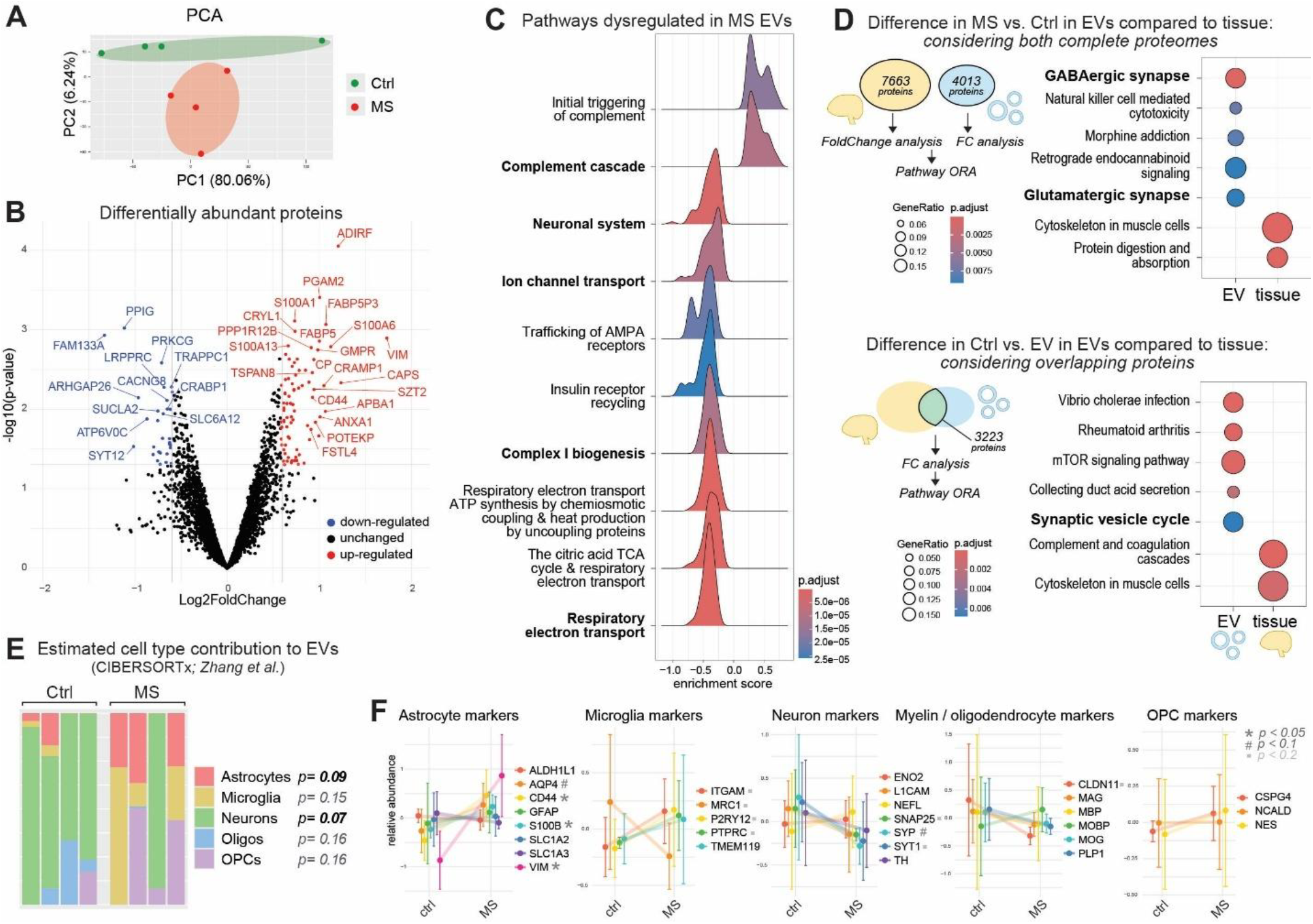
Proteomic changes in MS NAWM EVs compared to control WM EVs. **(A)** Principal component analysis using normalized abundances of proteins with MS EV samples in red and control (Ctrl) EV samples in green. **(B)** Volcano plots showing up- (red) and downregulated (blue) proteins with the following cutoffs: p-value < 0.05 and a < -1.5 or > 1.5-fold change. Proteins are shown as gene symbols. **(C)** Ridge plots of GSEA showing the enriched REACTOME pathways in MS compared to control EVs (same analysis for tissue shown in supplementary Fig. S2A). **(D)** Dot plots showing the pathways dysregulated in MS overrepresented in EVs compared to tissue, using two approaches: either including all proteins detected in the two proteomics datasets or including only the proteins present/overlapping in both datasets. **(E)** Relative CNS cell contribution to the WM EV pool in MS and control EVs estimated using CIBERSORTx cell deconvolution analysis. Each bar represents a sample, and each color a cell type (a comparison with the changes in the tissue is shown in Fig. S2). Statistical significance was determined using a Student’s t-test. **(F)** Graphs showing mean relative abundance of selected astrocyte, microglial, neuronal, myelin/oligodendrocyte, and OPC markers in control and MS EVs (mean protein abundance ± SD; n = 4 samples). Statistical significance indicated is from the limma differential abundance analysis, which is based on linear models and empirical Bayes methods.

These changes were confirmed in pathway analysis: pathways related to homeostatic neuronal and mitochondrial functions were downregulated, and inflammatory pathways, particularly the complement cascade, were upregulated in MS EVs compared to control EVs (Fig. 2C). Similar pathways were also dysregulated in the corresponding tissue (supplementary Fig. S2A-B). To identify which pathways are more strongly dysregulated in EVs compared to tissue, we carried out pathway overrepresentation analysis comparing the pathways dysregulated between MS vs. Ctrl tissue in EVs and tissue. This analysis was carried out using the entire proteome for the EV and tissue datasets and repeated only using the overlap of the two datasets. While MS-induced changes in neuronal/synaptic pathways were more pronounced in EVs, MS-induced changes in cytoskeletal pathways and the complement cascade were more pronounced in the tissue (Fig. 2D).

To determine if the cells contributing to the EV pool change in MS, we carried out CNS cell deconvolution analyses on the EV proteomics data using CIBERSORTx *(16–18)* and BRETIGEA *(19)*. In both analyses, a trend of increasing astrocyte and decreasing neuronal contributions (driven mainly by synaptic proteins) was observed, but this did not reach significance in either analysis (Fig. 2E and supplementary Fig. S2C). The trends observed in the BRETIGEA deconvolution analyses aligned with the results in the CIBERSORTx analysis (Fig. S2C) and with changes in commonly used CNS cell markers (Fig. 2F). Astrocyte cell markers for reactive subsets (VIM, CD44, S100B) were particularly strongly upregulated in MS EVs.

### Pathways related to synaptic signaling, mitochondrial function, complement activation, and aging are dysregulated in MS tissue EVs compared to controls

Neuronal system, mitochondria/respiratory electron transport chain, and complement cascade-related pathways were among the top dysregulated pathways in MS tissue-derived EVs. All three of these pathways have been shown to play a role in MS disease progression and white matter pathology. Therefore, we next characterized which proteins contribute to the changes in these pathways.

The most strongly dysregulated proteins within the neuronal system pathway were synaptic proteins (Fig. 3A). Consistent with our earlier analyses, these changes were more pronounced in EVs than in the corresponding tissue samples (supplementary Fig. 3A). To investigate whether specific subsets of synaptic proteins were particularly affected, we categorized proteins as uniquely pre- or postsynaptic by identifying those annotated with Cellular Component Gene Ontology (CC GO) terms containing either “presynaptic” but not “postsynaptic,” or “postsynaptic” but not “presynaptic.” Both pre- and postsynaptic proteins were enriched at the bottom of the ranked list of EV proteins, ordered by decreasing fold-change, suggesting they are downregulated. In direct comparison, presynaptic proteins exhibited a lower (more negative) enrichment score compared to postsynaptic proteins (NES pre: -1.94; post: 1.70), indicating more pronounced downregulation of presynaptic proteins (Fig. 3B). To compare changes in inhibitory and excitatory synaptic proteins, we identified these proteins by searching the GO terms for either “GABA-ergic” and “inhibitory” or “glutamatergic” and “excitatory”. The enrichment analysis revealed downregulation of both inhibitory and excitatory synaptic proteins, with inhibitory proteins having a lower enrichment score (NES inhib.: -2.19; exci.: -1.53), suggesting an overall more pronounced downregulation compared to excitatory synaptic proteins (Fig. 3B). These findings were further confirmed using the SynGO synapse function enrichment tool *(20)*. There was an overall enrichment of synaptic proteins, with a stronger enrichment of presynaptic proteins compared to postsynaptic proteins among the differentially regulated proteins in MS EVs (Fig. 3C).

**Fig. 3.**
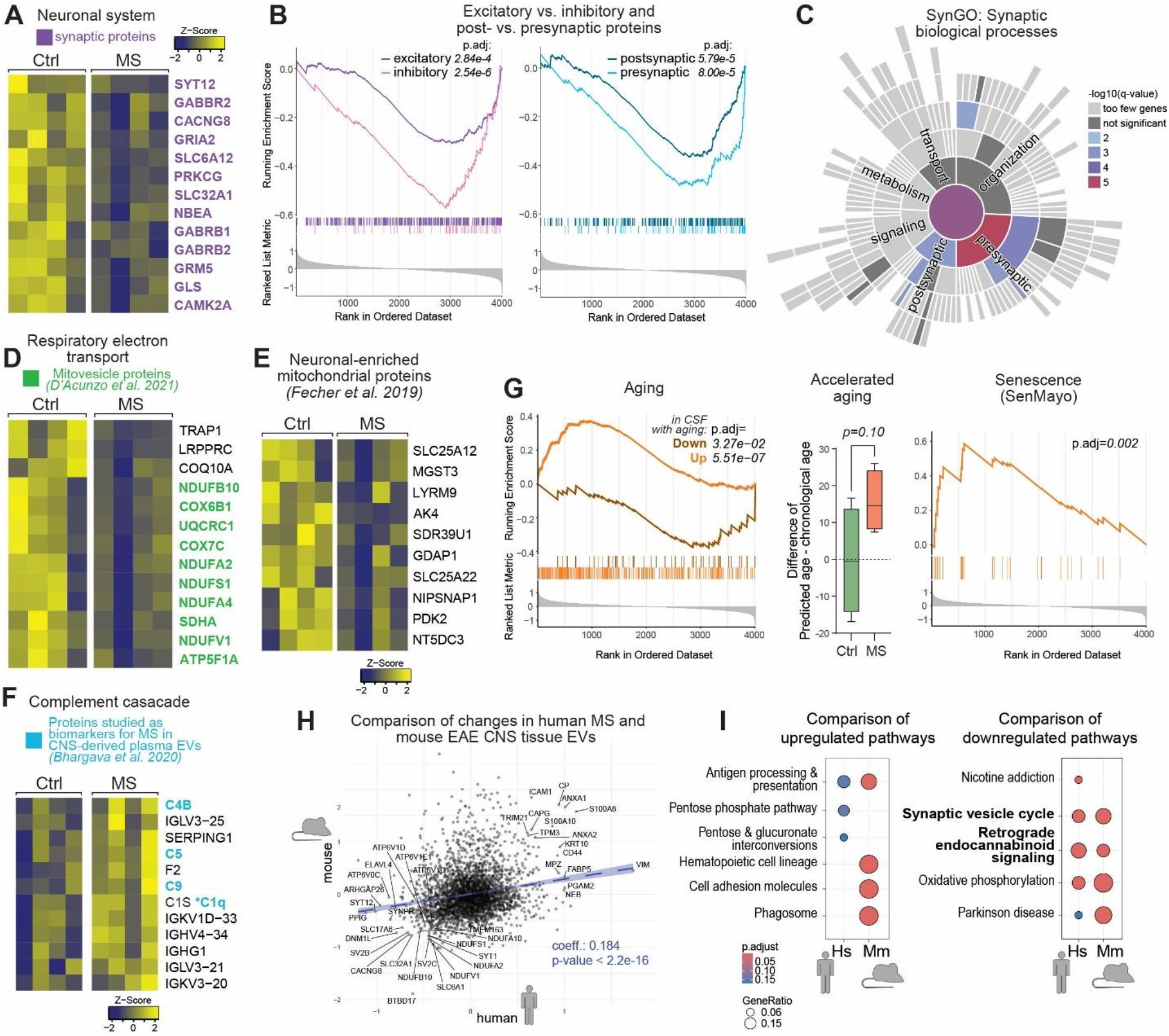
Pathways related to synaptic and mitochondrial functions, complement activation, and aging are dysregulated in MS NAWM EVs, and changes in EAE CNS EVs reflect changes in human MS EVs. **(A)** Heatmap showing the relative abundance of the top dysregulated proteins from the “Neuronal System” pathway with all synaptic proteins highlighted in purple. **(B)** GSEA for excitatory/ inhibitory and post-/presynaptic protein sets (based on GO terms) showing enrichment in MS EVs compared to control EVs. **(C)** Sunburst plot of SynGO *(20)* pathway analysis. **(D)** Heatmap showing the relative abundance of the top dysregulated proteins from the “Respiratory electron transport” pathway, with all proteins that have also been reported to be present in mitovesicles *(26)* highlighted in green. **(E)** Heatmap showing the relative abundance of neuronal-specific mitochondrial proteins *(28)* (only the top dysregulated proteins are shown). **(F)** Heatmap showing the relative abundance of the top dysregulated proteins from the “Complement cascade” pathway with all proteins that have been studied as EV biomarkers *(29, 30)* in MS highlighted in blue. **(G)** GSEA of EV samples for a set of proteins downregulated (brown) and upregulated (orange) in the CSF with aging on the left and for a set of senescence-related proteins (orange) on the right *(21, 22)*. The same analysis was carried out for the tissue samples (see supplementary Fig. S3). The box and whiskers plot in the center shows the difference in predicted (using a CSF proteomic clock tool *(22)*) and chronological age. For this analysis statistical significance was determined using a Student’s t-test. **(H)** Scatter plot showing the correlation of log2-fold-changes in proteins in human MS brain-derived EVs and mouse EAE spinal cord-derived EVs *(6)*. **(I)** Dot plots showing the pathways up-/downregulated in MS/EAE EVs overrepresented in humans compared to mice. All proteins are shown as gene symbols.

The top dysregulated proteins in the respiratory electron transport pathway were all mitochondrial proteins, many of which, interestingly, have previously been shown to be present in mitovesicles *(26)* (Fig. 3D; top dysregulated mitovesicle proteins are shown in supplementary Fig. S3B). Axons in MS NAWM contain fewer mitochondria (unless undergoing remyelination) and show signs of mitochondrial dysfunction *(27)*. Based on this, we hypothesized that the reduction of mitochondrial proteins may be partially driven by changes in neuron-derived EVs. Our dataset supports this hypothesis, revealing a reduction in neuronal-specific mitochondrial proteins *(28)* in MS EVs (Fig. 3E; astrocyte-specific mitochondrial proteins showed no consistent changes Fig. S2C).

The complement cascade pathway upregulation in MS EVs was driven by both initiators (immunoglobulins), components (C1s, C4B, C5, C9), and inhibitors (SERPING1) of the complement cascade (Fig. 3F), some of which have been studied as EV-based biomarkers in MS *(29, 30)*.

All three pathways discussed above are associated with brain aging. Therefore, we next tested for enrichment of aging-related proteins in MS EVs. We found that proteins known to be up-or downregulated in the CSF with aging *(22)* are enriched at the upper and lower ends, respectively, of the ranked list of EV proteins, ordered by decreasing fold-change. This suggests that these aging-related proteins are dysregulated in MS EVs (Fig. 3G). This finding was further corroborated by calculating the predicted age of MS EVs samples using a clock tool based on the CSF proteomics dataset mentioned above *(22)*. While MS EVs exhibited signs of accelerated brain aging compared to control EVs, the difference did not reach statistical significance given our sample size. Using the SenMayo dataset *(21)*, we further found that proteins associated with senescence, a hallmark of aging, are also upregulated in MS EVs compared to control EVs.

Finally, we assessed whether the proteomic changes observed in human MS-derived EVs were conserved in the mouse EAE model. Indeed, we found a significant correlation in the log2-fold changes of EV proteins between MS and EAE. Comparison of the overrepresented pathways among dysregulated proteins revealed that both synaptic and mitochondrial pathways were consistently downregulated in EVs from MS and EAE. In contrast, there was less overlap in the upregulated pathways. Specifically, inflammatory pathways, including antigen presentation and phagocytosis, were overrepresented in EAE EVs but were either unchanged or only slightly changed in MS EVs. This might be due to differences in the tissues analyzed in mice and humans: while EAE EVs were separated from whole spinal cords, likely containing active demyelinating lesions, human MS EVs were separated from NAWM, which lacks lesional tissue.

## DISCUSSION

Overall, there were no significant differences in the median size, yield, and morphology in MS white matter EVs compared to controls. The slight shift towards larger EVs observed in the size distribution histogram may suggest alterations in the mechanisms of EV release or changes in the cell of origin in MS. This might be linked to the inflammation in NAWM as the size and number of EVs released by some glial cells have been shown to increase under inflammatory conditions *(31–33)*.

Our cellular deconvolution analyses revealed a trend toward increased astrocytic and decreased neuronal contributions to the EV pool in MS compared to controls. This shift may reflect changes in the cellular composition of the NAWM, e.g., more astrocytes and/or neuronal loss, or altered EV production by these cell types. Supporting the former, tissue deconvolution analysis revealed similar MS-associated shifts in predicted cell type proportions, suggesting that EV changes may partially mirror tissue composition. However, increased EV production by reactive astrocytes may also contribute. In vitro studies have shown that reactive astrocytes release more EVs under inflammatory conditions *(32, 33)*, which aligns with our observation of upregulated reactive astrocyte markers, such as CD44 and Vimentin, in MS-derived EVs.

In MS-derived EVs, synaptic signaling and mitochondrial oxidative phosphorylation pathways were downregulated, while complement cascade-related pathways were upregulated compared to controls. These changes are particularly interesting given that these pathways are key features of MS pathology. MS involves synaptic pathology, even in the absence of demyelination and axonal loss *(34)*, highlighting its relevance as an independent pathological feature of MS. Under homeostatic conditions, neurons release EVs containing synaptic proteins, which support synaptic plasticity and provide trophic signals to stressed neurons *(35)* (reviewed in more detail elsewhere *(36)*). The reduced synaptic cargo in MS EVs may therefore not only reflect ongoing synaptic pathology but could also perpetuate synaptic loss and neuronal stress in MS. Notably, the more pronounced loss of inhibitory and presynaptic proteins aligns with the pattern of synaptic loss observed in MS tissue and animal models of MS *(37–39)*. However, what was surprising was that these changes in synaptic proteins were observed in NAWM EVs, despite synaptic pathology in MS primarily being described in the gray matter. One possible explanation is that some of the EVs detected in NAWM originate from other brain regions, including the gray matter. Alternatively, these findings may suggest that synaptic pathology also occurs within the white matter itself. Although classical synapses are rare in the white matter, axon-glial synapses have been described in white matter *(40–42)*. These synapses have been shown to contain both inhibitory and excitatory synaptic proteins and regulate remyelination. Disruption of these synapses in MS – perhaps due to Wallerian degeneration, reduced proximal input secondary to synaptic pathology in the GM, or compartmentalized inflammation – could also explain the changes in synaptic proteins in MS NAWM EVs.

The MS-associated synaptic pathology and neurodegeneration described above are linked to mitochondrial dysfunction *(43)*, which we also saw in MS EVs. There was an overall loss of mitochondrial proteins with a marked downregulation of complex I-associated proteins. This mitochondrial dysfunction, including the loss of neuronal complex I, is well described in neurons in the gray matter *(43)*. In white matter, especially NAWM, the patterns are less well understood, in part because most studies in MS white matter focus on changes in MS lesions, only including NAWM as internal control tissue. In close proximity to MS lesions, axons (demyelinated and remyelinated), microglia, and astrocytes have increased mitochondrial density and complex I activity, which sustains neuroinflammation *(27, 44, 45)*. The MS NAWM, on the other hand, has fewer mitochondria in myelinated axons and shows lower levels of the neuronal mitochondrial metabolite N-acetylaspartate in progressive disease stages compared to healthy control WM *(27, 46)*. The reduced mitochondrial proteins in MS may therefore reflect mitochondrial dysfunction in NAWM axons, especially since we saw a pronounced change in neuron-specific mitochondrial proteins. However, besides merely reflecting disease pathology, the loss of mitochondrial proteins in EVs might also contribute to MS disease processes. EVs can shuttle mitochondrial proteins and even entire functional mitochondria *(26, 47, 48)*. Mitochondrial cargo in CNS cell EVs, including neuronal progenitor and astrocyte EVs, has been shown to reduce the inflammatory activation of mononuclear phagocytes and promote neuronal cell survival *(47, 48)*. Furthermore, depolarized mitochondria contributing to intracellular oxidative stress can be released in EVs and taken up by recipient cells, which then restore the mitochondria’s metabolic functions *(49)*. The reduced mitochondrial cargo in MS EVs is indicative that MS tissue lacks the metabolic reserve/ regenerative capacity these EVs offer. Hence, the loss might perpetuate mitochondrial damage and downstream oxidative stress in MS. Enhancing EV-mediated mitochondrial protein transfer could then be a potential therapeutic strategy in MS.

Complement and triggering of the complement cascade were among the top up-regulated pathways. In MS NAWM pathology, CNS complement activation plays diverging roles, promoting debris clearance through phagocytosis but also contributing to the compartmentalized inflammation and tissue damage when activated excessively or continuously (reviewed in more detail elsewhere *(50)*). The complement cascade is thought to mediate many of these effects through extracellular signaling of the soluble complement components. However, there is increasing evidence that the complement components can be activated and can execute their functions intracellularly as part of the complosome *(51, 52)*. In this context, our findings of changed complement pathway-related proteins in MS EVs are particularly interesting because they suggest that, besides its extracellular and intracellular functions, complement might also play a role in EVs. There is some evidence for this in C3-containg fibroblast-derived EVs, which have been shown to induce inflammatory activation and metabolic reprogramming when taken up by macrophages *(53)*. Further microglial EVs have been shown to bind C1q on their surface, shuttling it to synapses to promote synaptic pruning *(54)*. Should a similar mechanism be taking place in MS, the upregulation of complement proteins in EVs could contribute to CNS compartmentalized inflammation and (excessive and/or protective) synaptic pruning in MS. Whether the complement related proteins detected in EVs are within EVs or bound to their surface externally or possibly even part of the EV biocorona is unclear and should be subject of future investigations.

Complement deposition, synaptic pathology, and mitochondrial dysfunction are all associated with brain aging (reviewed in more detail here *(55)*). We therefore hypothesized that EVs might show signs of accelerated brain aging, which we saw trending. Further, senescence-associated proteins were upregulated in MS EVs, including ANXA1 and S100 proteins, which have previously been shown to be markers of senescent EVs *(56)*. Senescent EVs can contribute to inflammation (reviewed in more detail *(57)*) and more recently have also been suggested as mediators of paracrine senescence independent of the SASP *(58, 59)*.

Interestingly, several of our findings in MS tissue EVs align with what has been shown in EVs derived from the CSF and blood of MS patients*(29, 30, 60)*. We found a trend of increased astrocytic and neuronal EV contribution to the tissue EV pool in MS compared to controls. In line with this, astrocytic EVs (identified either as GLAST+ EVs or EAAT2+ EVs) have been shown to be increased both in blood and CSF *(29, 61)* and neuronal EV particle counts and the expression of neuronal markers are decreased in circulating EVs from MS patients compared to healthy controls *(30, 62)*. Though this is not consistent among all studies *(29)*, which may be due to differences in the cellular markers used, EV particle quantification methods, and patient cohort characteristics across studies. Besides the cellular contribution, the loss in synaptic proteins we observed in MS tissue EVs also aligned with findings in circulating EVs from MS patients, where synaptic protein levels are reduced in neuronal EVs compared to healthy controls, and the rate of synaptic protein loss is associated with brain and retinal atrophy *(29, 63)*. Lastly, the increase in complement-related proteins we saw in MS tissue EVs has also been reported in both plasma and CSF EVs. Circulating astrocyte-enriched EVs from MS patients contain higher levels of several complement proteins, including those identified in this study, and complement pathways are enriched in RRMS CSF EVs compared to control EVs and whole CSF *(29, 60)*. Our study provides compelling evidence that the MS-related changes in EVs in the brain may be detectable in circulation, which is an exciting prospect because CNS-derived EVs in the circulation might provide a unique opportunity to monitor progressive disease stages, which are driven by CNS-internal processes, rather than peripheral immune activation. To better characterize the overlap of CNS tissue, CSF, and circulating EVs, it will, however, additionally be necessary to carry out studies with samples from all three compartments. Since it is hard to come by these samples in humans, these experiments could also be carried out animals models of MS, in which we found similar changes in the CNS EV proteome *(6)*.

The primary limitation of this study is the small sample size, which restricts the statistical power and generalizability. Future studies should aim to expand the sample size, allowing for the breakdown of samples according to MS subtypes or possibly even novel glia biology-driven MS subclassifications *(64)*. Additionally, comparisons between different lesion types and NAWM within the same patient would provide deeper insights into how EVs might be impacted by and possibly even contribute to the distinct molecular processes taking place in these regions. The EV separation method used, while effective for separating EVs from human brain tissue, does not capture the full range of particles and, like every separation method, has its unique set of disadvantages (reviewed in more detail here *(65)*). Complementary studies using different separating methods may yield slightly different results and further enrich our understanding of EVs in MS. Moreover, the bioinformatics tools used in this study, such as the cell deconvolution analyses, the pathway analyses, and the aging clock tool, were not specifically developed for EV proteomics and may therefore not fully capture changes in EV-specific pathways or cellular markers.

In conclusion, this study found no significant differences in WM CNS-tissue EV median size, yield, or morphology in MS compared to controls, though there was a slight shift towards larger EVs in MS. There was trend in increased astrocytic EVs and decreased neuronal EVs, accompanied by alterations in pathways related to synaptic signaling, mitochondrial function, complement activation, and aging in MS tissue EVs compared to controls. These findings are consistent with MS pathology and suggest that EVs may play a role in mediating these disease processes. The similarities with changes seen in circulating MS EVs confirm that circulating CNS-derived EVs are a promising platform to monitor CNS disease progression. Our study provides a novel resource to select interesting biomarker candidates.

## Geolocation

North America

## Acknowledgements

We thank Matthew Smith for his valuable insights and helpful discussions regarding the data analysis.

## Supplementary material

Supplementary Figures S1-S3; Supplementary Tables 1-3; and Supplementary Method File.

