## Supplementary Figures and Methods for "Extracellular Vesicles from Multiple Sclerosis White Matter Exhibit Synaptic, Mitochondrial, Complement, and Aging-related Pathway Dysregulation"

#### Supplementary Tables (in separate excel file)

- Supplementary table 1: EV proteomics data
- Supplementary table 2: tissue proteomics data
- Supplementary table 3: Extended data on patients included in this study

#### Supplementary Figures

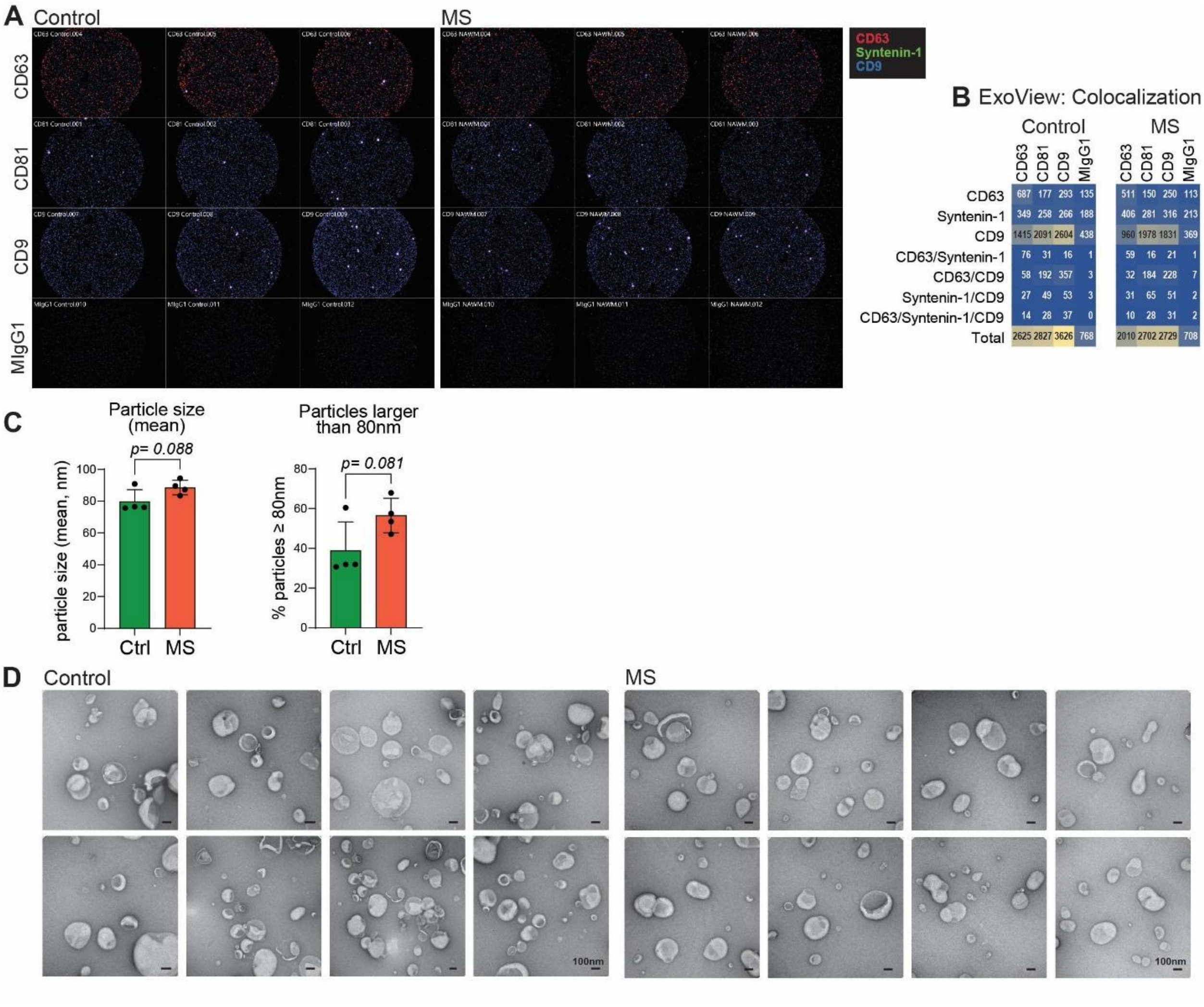


**Supplementary Fig. S1: Extended data for EV characterization. (A)** SP-IRIS images with rows representing the markers used to capture EVs and colors representing the secondary detection antibodies (CD63 in red, Syntenin-1 in green, and CD9 in blue). **(B)** Heatmaps showing the number of particles with colocalization of EV markers, where columns represent the markers used to capture EVs, and rows represent the fluorescent signals of the detection antibodies and their colocalization. **(C)** Bar graphs showing the mean particle size (the main figure shows median size) and percentage of particles larger than 80nm ± SD measured using nanoflow cytometry (n = 4 samples for MS NAWM and control [Ctrl] WM EV samples each). Statistical significance was determined using a Student's t-test (ns = not significant). **(D)** Electron microscopy images of negatively stained EVs from control WM and MS NAWM (at a higher resolution than in Fig 1).

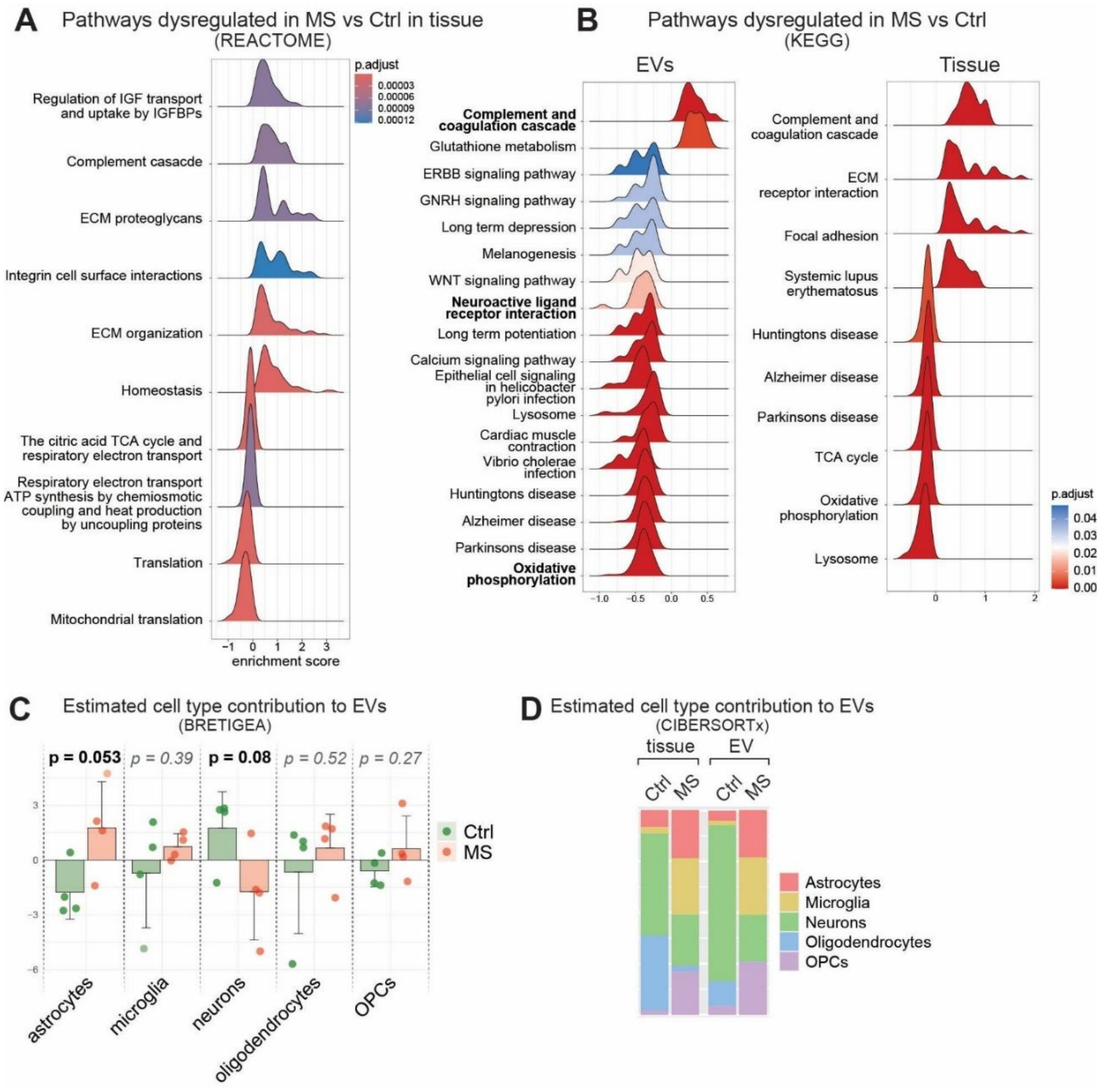


**Supplementary Fig. S2: Extended pathway and CNS cell deconvolution analyses. (A)** Ridge plots of GSEA showing the enriched REACTOME pathways in MS compared to control tissue (same analysis for EVS shown in supplementary Fig. 2). **(B)** Ridge plots of GSEA showing the enriched KEGG pathways in MS compared to control EVs (left) and tissue (right). **(C)** Bar plot of the BRETIGEA deconvolution score for each cell type in control (green) and MS (red) EVs. Statistical significance was determined using a Student's t-test. **(D)** Relative CNS cell contribution to the protein pool in MS and control tissue and EVs estimated using CIBERSORTx cell deconvolution analysis. Each bar represents a condition (shown is the mean contribution of each cell type), and each color represents a cell type.

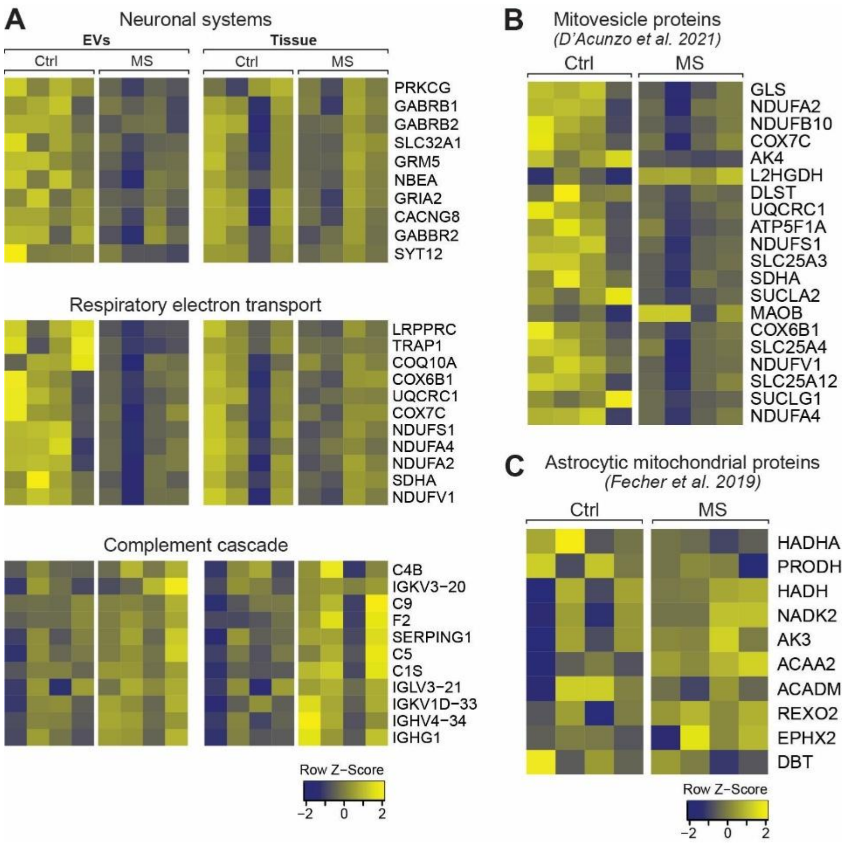


**Supplementary Fig. S3: Extended plots for pathways/proteins of interest and aging analysis in tissue. (A)** Heatmap showing the relative abundance of the top dysregulated proteins in both EVs and tissue from the “Neuronal System”, “Respiratory electron transport” and “Complement cascade” pathways. **(B)** Heatmap showing the top dysregulated mitovesicle proteins in EVs of which most (all but two) are downregulated. **(C)** Heatmap showing the top dysregulated astrocyte-specific mitochondrial proteins, which, in contrast to the neuronal-specific mitochondrial proteins (shown in Fig. 3E), do not show a consistent pattern of change.

### Supplementary methods

All relevant references for this section have already been cited in the methods section of the manuscript.

#### Separation of Extracellular Vesicles from Brain Tissue

Brain-derived extracellular vesicles (EVs) were separated according to a previously published protocol *(4)*. Briefly, frozen human tissue from normal-appearing white matter of MS and control samples was gently sliced into 2–4 mm pieces on dry ice. A small (~50 mg) piece of tissue was stored at –80 °C for later protein extraction from brain homogenate (BH). Chopped tissue (~400 mg) was transferred to a 15 mL tube containing 75 U/mL Collagenase Type III (Worthington #CLS-3) in Hibernate-E medium (Thermo #A12476-01) at a ratio of 800 µL per 100 mg of tissue. Samples were incubated at 37 °C for 20 minutes. After incubation, the mixtures were placed on ice and supplemented with protease and phosphatase inhibitors. Samples were then subjected to differential centrifugation: first at 300×g for 10 minutes to remove debris, followed by 2,000×g for 15 minutes at 4 °C. The supernatant was filtered through a 0.22 µm filter (Millex GS), then centrifuged at 10,000×g (10K) for 30 minutes at 4 °C. The resulting EV-containing supernatant was concentrated using a 100 kDa molecular weight cutoff concentrator (Amicon Ultra) to a final volume of 700 µL. Concentrated samples were loaded onto a rehydrated qEV original 70 nm column (IZON), and EV fractions were collected per the manufacturer's instructions. Pooled EV fractions were centrifuged at 110,000×g for 90 minutes at 4 °C. The supernatant was removed, and the pellet resuspended in 70 µL DPBS containing protease and phosphatase inhibitors. EVs were then aliquoted into 50 µL portions for proteomic analysis and 20 µL for characterization.

#### Transmission Electron Microscopy

A droplet of freshly thawed EV suspension was adsorbed onto glow-discharged, ultrathin carbon-coated 400 mesh copper grids (EMS CF400-CU-UL) for 2 minutes. The grids were then washed three times with TBS and once with distilled water. Negative staining was performed using 1% uranyl acetate (UA) in deionized water with 1% tylose. Samples were incubated with 4% UA for 40 seconds for contrast, blotted dry, and imaged using a Hitachi 7600 transmission electron microscope. Eight images were captured per pooled EV sample (consisting of all four samples for each condition).

#### EV Quantification by Nanoflow Cytometry

EV concentration and size distribution were measured using the NanoFCM Flow Nano Analyzer according to the manufacturer’s instructions. Calibration was performed separately for particle concentration (using 250 nm silica nanoparticles, NanoFCM #QS2503) and size (using a premixed cocktail of 68, 91, 113, and 155 nm nanoparticles, NanoFCM #516M-Exo). DPBS served as the blank. EVs were diluted 1:1000 in DPBS. Signal acquisition was performed for 1 minute at 1 kPa with an event rate of 1,500–10,000 events/min. Particle size and count were determined using calibration curves and side scatter intensity in NanoFCM Professional Suite v2.0.

#### Single-Particle Interferometric Reflectance Imaging Sensor (SP-IRIS)

EVs were phenotyped using the ExoView R100 scanner (NanoView Biosciences) and the EV-TETRA-C Tetraspanin Kit per the manufacturer’s protocol. Ten µL of EVs were diluted with 35 µL incubation buffer and loaded onto ExoView chips, then incubated at room temperature for 16 h. After four washes with incubation buffer (3 min each), chips were incubated with fluorescent antibodies against human CD63, CD9, and Syntenin-1 (1:1200 dilution in a 1:1 mix of incubation and blocking buffers). After further washing (IB, wash buffer, rinse buffer), chips were scanned using SP-IRIS and fluorescence detection. Data were analyzed using NanoViewer v2.8.10.

#### Sample Preparation and LC-MS/MS Analysis

EVs were dried under vacuum and reconstituted in 8 M urea with 50 mM triethylammonium bicarbonate (TEAB). Samples were reduced and alkylated with 10 mM TCEP and 40 mM CAA at room temperature for 1 hour, then digested with Lys-C (1:100) for 3 hours at 37 °C. After diluting urea to 2 M, samples were further digested with trypsin (1:50) overnight. Peptides were desalted using C18 StageTips and labeled with TMTpro 18-plex reagents. Labeled peptides were fractionated into 96 fractions by basic pH reversed-phase LC, concatenated into 24 pools, vacuum-dried, and reconstituted in 0.5% formic acid. Peptides were analyzed using an Orbitrap Fusion Lumos mass spectrometer coupled with an Ultimate 3000 RSLCnano system. MS1 scans (m/z 300–1800) and MS2 scans were acquired in data-dependent acquisition mode (Top Speed, 3 s cycle). MS1 and MS2 resolutions were 120,000 and 50,000, respectively. HCD was used for fragmentation with 35% NCE. The AGC targets were 1e6 (MS1) and 5e4 (MS2). Dynamic exclusion was 30 s. Internal lock mass calibration was enabled (m/z 445.12002).

**Database Search and Statistical Analysis**

MS/MS data were searched against the UniProt human database (Swiss-Prot + TrEMBL, Jan 2019 release) using MSFragger v3.4 within Proteome Discoverer v2.4. Fixed modifications included carbamidomethylation (+57.021 Da) and TMTpro tagging (+304.207 Da); methionine oxidation (+15.995 Da) was set as a variable modification. A maximum of two missed cleavages was allowed. Precursor and fragment tolerances were set at 10 ppm and 20 ppm, respectively. FDR filtering was set to 1% for both peptide and protein levels. Quantification was based on centroid reporter ions with a 20 ppm tolerance and S/N >10. Only unique and razor peptides were used. The coisolation threshold was 50%. Missing intensity values were imputed with the minimum value. Protein grouping followed strict parsimony, and proteins without unique peptides were removed.

**Sample Preparation and LC-MS/MS Analysis**

Sample preparation and liquid chromatography–tandem mass spectrometry (LC-MS/MS) analysis were performed as previously described, with minor modifications *(6)*. EVs were vacuum-dried and reconstituted in a lysis buffer containing 8 M urea and 50 mM triethylammonium bicarbonate (TEAB; both Sigma-Aldrich). Reduction and alkylation were carried out in the presence of 10 mM tris(2-carboxyethyl) phosphine (TCEP) and 40 mM 2-chloroacetamide (CAA; Sigma-Aldrich) at room temperature for 1 hour.

Proteins were digested with Lys-C (MS grade; Fujifilm Wako Pure Chemical Industries Co., Ltd.) at a 1:100 enzyme-to-protein ratio for 3 hours at 37 °C. Subsequently, the urea concentration was diluted to 2 M by adding three volumes of 50 mM TEAB. Trypsin (sequencing-grade modified; Promega) was then added at a 1:50 ratio, and samples were incubated overnight at 37 °C. Peptides were desalted using C18 StageTips (3M Empore) and labeled with TMTpro 18-plex reagents (Thermo Fisher Scientific) following sample randomization and per the manufacturer's instructions.

Labeled peptides were pre-fractionated using basic pH reversed-phase liquid chromatography into 96 fractions, which were concatenated into 24 pooled samples. These were vacuum-dried and reconstituted in 0.5% formic acid (FA; Thermo Fisher Scientific) for LC-MS/MS analysis. Peptides were analyzed on an Orbitrap Fusion Lumos Tribrid mass spectrometer coupled to an Ultimate 3000 RSLCnano system (both Thermo Fisher Scientific).

Peptides were first loaded onto a trap column (Acclaim PepMap 100 C18, 5 µm, 100 µm × 2 cm, nanoViper) at 8 µL/min and separated at 0.3 µL/min on an analytical column (Easy-Spray PepMap RSLC C18, 2 µm, 75 µm × 50 cm) using a 120-minute linear gradient increasing solvent B (0.1% FA in 95% acetonitrile) to 28%. The EASY-Spray source was operated at ~2.5 kV.

Mass spectrometry was performed in data-dependent acquisition (DDA) mode using a Top Speed method with a 3-second cycle time. MS1 scans were acquired from m/z 300–1,800 at 120,000 resolution; MS2 scans were acquired at 50,000 resolution using higher-energy collisional dissociation (HCD) at 35% normalized collision energy. The AGC targets were set to $1 \times 10^6$ (MS1) and $5 \times 10^4$ (MS2), with maximum ion injection times of 50 ms and 100 ms, respectively. The precursor isolation window was 1.6 m/z with a 0.4 m/z offset. Dynamic exclusion was enabled for 30 seconds, and singly charged ions were excluded. Internal calibration was performed using the lock mass (m/z 445.12002) from ambient air.

**Database Searches and Statistical Analysis**

MS/MS data were searched against the UniProt human protein database (Swiss-Prot and TrEMBL, January 2019 release), including common contaminants. Searches were conducted using the MSFragger v3.4 algorithm via the Proteome Discoverer software (v2.4.1.15; Thermo Scientific) *(7)*.

During MS2 preprocessing, the top ten peaks in each 100 m/z window were selected for database searching. Trypsin was set as the protease, allowing up to two missed cleavages. Fixed modifications included carbamidomethylation of cysteine (+57.021 Da) and TMTpro labeling on lysine residues and peptide N-termini (+304.207 Da). Methionine oxidation (+15.995 Da) was specified as a variable modification. A minimum peptide length of six amino acids was required.

Precursor (MS1) and fragment (MS2) mass tolerances were set at 10 ppm and 20 ppm, respectively. False discovery rate (FDR) filtering was applied at 1% for both peptide and protein levels using the Percolator and Protein FDR Validator nodes. Protein quantification was performed using the most confident centroid method with a reporter ion tolerance of 20 ppm. Peptide quantification included both unique and razor peptides, and protein groups were defined by peptide uniqueness. The coisolation threshold was set at 50%.

Reporter ion intensities were based on signal-to-noise (S/N) ratios, with an average S/N threshold of 10. Missing intensity values were imputed with the dataset minimum. Isobaric tag correction and normalization were disabled. Protein grouping followed a strict parsimony approach, with proteins lacking unique peptides excluded. Final protein groups were generated by selecting peptide-spectrum matches (PSMs) with the highest number of unambiguous peptides and summing reporter ion intensities for quantification.

**Further Bioinformatic and Statistical Analysis**

Subsequent analyses were conducted in R (v4.3.1). UniProt IDs were mapped to Entrez IDs and gene symbols using the AnnotationDbi package and the org.Hs.eg.db annotation database. Principal component analysis (PCA) was performed using the stats package and visualized with the autoplot function from ggplot2. Proteins with missing values in any sample were excluded from downstream analyses. Data were normalized using median centering and log2-transformed.

Differential abundance analysis was carried out using the limma package, which fits a linear model (analysis of variance, ANOVA) and applies an empirical Bayes moderation to the test statistics. Volcano plots and heatmaps were generated using ggplot2, ggrepel, and the heatmap.2 function from the gplots package.

Pathway analyses included over-representation analysis (ORA) and gene set enrichment analysis (GSEA), performed using the ClusterProfiler package. Gene sets tested included Gene Ontology terms, KEGG pathways, and Reactome pathways. Results were visualized using enrichplot and ggplot2.

To compare cellular component GO term enrichment in tissue versus EV samples, ORA was performed using the CompareCluster function on the top 10% most abundant proteins from each dataset. To assess pathway alterations in MS, ORA was conducted using either (1) the top dysregulated proteins in each dataset (tissue and EVs) or (2) the top dysregulated proteins shared between the two datasets.

Cell-type deconvolution was conducted using the CIBERSORTx online platform, employing a glial reference matrix derived from immunopanned human brain cell RNA-seq data. Observed trends were validated using the BRETIGEA package with singular value decomposition and 400 marker genes per cell type. Statistical testing of fold changes in estimated cell populations was performed using limma.

To further characterize changes in synaptic subtypes, cellular compartment GO terms associated with subsets of interest were queried, and resulting gene sets were subjected to GSEA. Presynaptic and postsynaptic protein changes were assessed by selecting GO terms containing either “presynaptic” but not “postsynaptic,” or vice versa. Excitatory and inhibitory synaptic proteins were analyzed using terms that included (1) both “GABAergic” and “inhibitory,” or (2) both “glutamatergic” and “excitatory.” SynGO-based functional pathway analysis and sunburst visualization were performed using the SynGO online platform.

To investigate whether MS EV proteomes exhibited signs of accelerated aging, GSEA was performed using aging-related gene sets from cerebrospinal fluid (CSF) proteomic datasets. Additionally, a CSF-based proteomic aging clock was used to calculate the difference between proteomic and chronological age in MS versus control samples.

EV proteomic changes observed in MS were compared to those detected in a chronic stage of experimental autoimmune encephalomyelitis. Log2-fold changes of overlapping proteins were correlated between species. ORA using CompareCluster was used to identify commonly or differentially regulated pathway clusters between MS and EAE. Because the mouse dataset exhibited lower variability (i.e., smaller p-values), the top 150 up- and downregulated proteins were selected for ORA.
